# Improved real-time influenza surveillance using Internet search data in eight Latin American countries

**DOI:** 10.1101/418475

**Authors:** Leonardo Clemente, Fred Lu, Mauricio Santillana

## Abstract

A real-time methodology for monitoring flu activity in middle income countries that is simultaneously accurate and generalizable has not yet been presented. We demonstrate here that a self-correcting machine learning method leveraging Internet-based search activity produces reliable and timely flu estimates in multiple Latin American countries.

## Introduction

With the highest mortality of any respiratory infectious disease in the young and elderly in Latin America, influenza poses significant health and economic challenges to low and middle income countries in the region [1]. The World Health Organization (WHO) maintains a healthcare-based disease surveillance system that collects information on flu activity from local ministries of health around the world. Unfortunately, these reports have a common delay of at least a week in Latin America, limiting the ability for a timely response to unexpected epidemic outbreaks. Reliable surveillance systems that monitor flu activity in real time in this region would help public health institutions deploy timely vaccination campaigns and optimally allocate resources during epidemic outbreaks. Multiple research teams have proposed complementary methods to estimate and forecast flu activity in real-time, in data-rich countries such as the US, using techniques ranging from statistical [2, 3] to mechanistic [4, 5], and incorporating a variety of data sources such as Internet search information, flu-related Twitter microblogs [6, 7], crowd-sourced flu surveillance [8, 9], clinician search activity [10], electronic health records [11], and Wikipedia access [12, 13], as summarized in [14]. However, a reliable system that leverages Internet search activity to monitor flu activity in multiple developing nations is not yet available.

An early large-scale implementation of real-time disease surveillance started in 2008 with Google Flu Trends (GFT), an online tool that used Google search activity to produce flu activity estimates in multiple locations around the world [15]. While GFT was initially perceived as a technological innovation, its large prediction errors during the 2009 H1N1 flu pandemic and the 2013 flu season in the U.S. raised methodological concerns from multiple researchers [16-18]. A recent study by Pollet showed that GFT flu estimates in Latin America had yielded poor results [19].

The discontinuation of GFT in 2015 led many to believe that Internet search trends were too noisy to track disease activity, a problem exacerbated in developing countries with limited Internet access. However, recent research has shown that robust and dynamically self-correcting machine learning methodologies can extract meaningful signals from real-time search activity to track Zika and Dengue activity in low to middle income countries around the world [11, 20, 21]. We apply lessons learned from these studies and successfully extend a state-of-the-art modeling approach for flu surveillance to eight Latin American countries.

## Data and Methods

We built a predictive methodology that estimates suspected flu activity in near real-time, as reported by FluNet, an online surveillance tool maintained by the WHO. FluNet collects and aggregates multiple indicators of flu activity at the country level. For this study, we selected the number of processed specimens (NPS) as ground truth. Since these specimens are taken from patients with flu-like symptoms and then sent to a laboratory for testing, we interpret them as an indicator of suspected flu activity in the population. Weekly aggregated NPS reports were collected from Jan. 5, 2009 to Dec. 25, 2016 for Argentina, Bolivia, Brazil, Chile, Mexico, Paraguay, Peru and Uruguay.

Given their near-real time availability, we selected influenza-related internet search activity to be used in our models as proxies for flu activity. We extracted a set of flu-related search term trends, individually for each country, collecting a total of 285 Spanish terms and 96 Portuguese terms. These search terms were identified using Google Correlate in countries where this tool was available. See Supplementary Materials for further description of this process.

We extended ARGO, a methodology originally conceived and tested to track flu activity in the U.S. in multiple spatial scales, as a way to produce retrospective and strictly out-of-sample flu estimates individually for each country [20, 22]. This methodology is based on a multivariable regularized linear model that is dynamically recalibrated every week as new flu activity information becomes available. Besides online search information, ARGO incorporates short-term and seasonal historical flu information to improve the accuracy of predictions and mitigate the undesired effect of spikes in search activity (induced perhaps by “panic” in the population during potential health threats reported by the news). More details on this approach can be found in [22].

Given a weekly as-yet-unseen NPS report to estimate, we used historical NPS and Google Trends information from the previous most recent 2 years (104 weeks) of data to calibrate ARGO and predict the given week’s NPS report. To assess ARGO’s predictive power, we built autoregressive models separately for each country (named AR52 throughout this paper) that only use historical flu activity from the 52 weeks prior to predictions, and generated retrospective out-of-sample estimates over the same time period. All models were built using the glmnet package on MATLAB version 2014a [3, 23].

To compare the predictive ability of ARGO and AR52, we calculated Person correlations and the root mean square error (RMSE) between model predictions and the subsequently observed suspected flu cases. The added value of using Google search activity as a predictor was tested via an efficiency metric [22] that quantifies the improvement of ARGO over a simple autoregressive model. This efficiency metric is calculated as the ratio between AR52’s and ARGO’s mean square errors. 90% confidence intervals for the efficiency metric were generated using the stationary block bootstrap method [24].

## Results

Retrospective out-of-sample estimates of flu activity were produced, for each of the 8 countries, from Jan. 1, 2012 to Dec. 25, 2016 and compared to the FluNet reported suspected cases (NPS). Brazil’s NPS data was only available until October 9, 2016. Note that due to FluNet’s reporting delays, our models, which rely on past available values of FluNet and current Internet search activity, estimate current flu activity at least one week ahead of official reports.

In Figure 1, we show our real-time flu estimates and the subsequently observed suspected flu cases for each country. For context, historical GFT values (scaled to be displayed alongside with NPS values) and autoregressive estimates are also shown. Our models (ARGO and AR52) accurately predict NPS values in each country. GFT shows consistently large discrepancies when compared to the observed values, consistent with the findings reported by [19].

**Figure 1.**
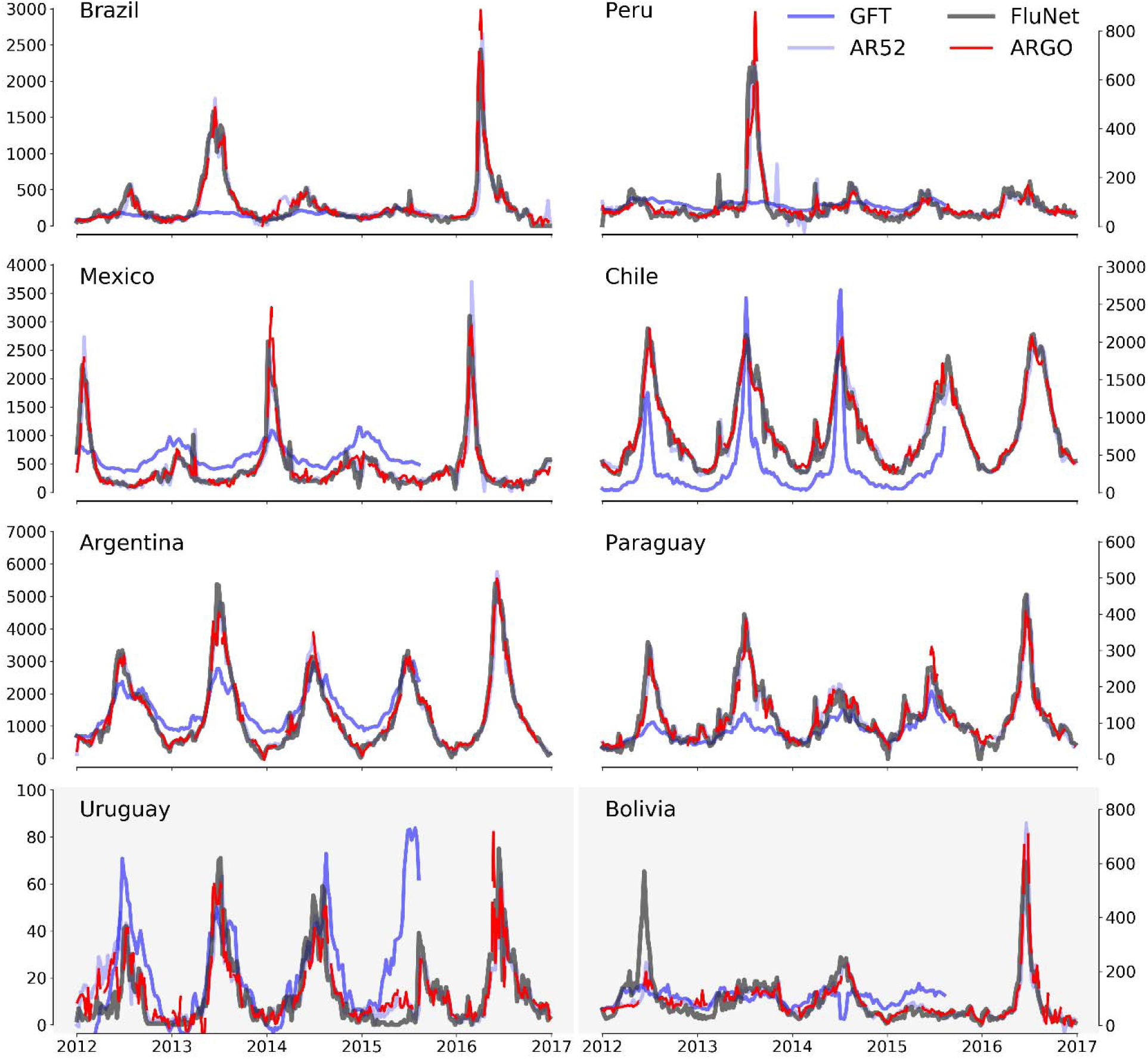
Graphical representation of the number of processed specimens (NPS) as reported by WHO’s Flunet (black), along with the NPS estimates generated by ARGO (red), AR (light blue) and Google Flu Trends (blue), over the whole study period of 2012-01-01 to 2016-12-25.

**Figure 2.**
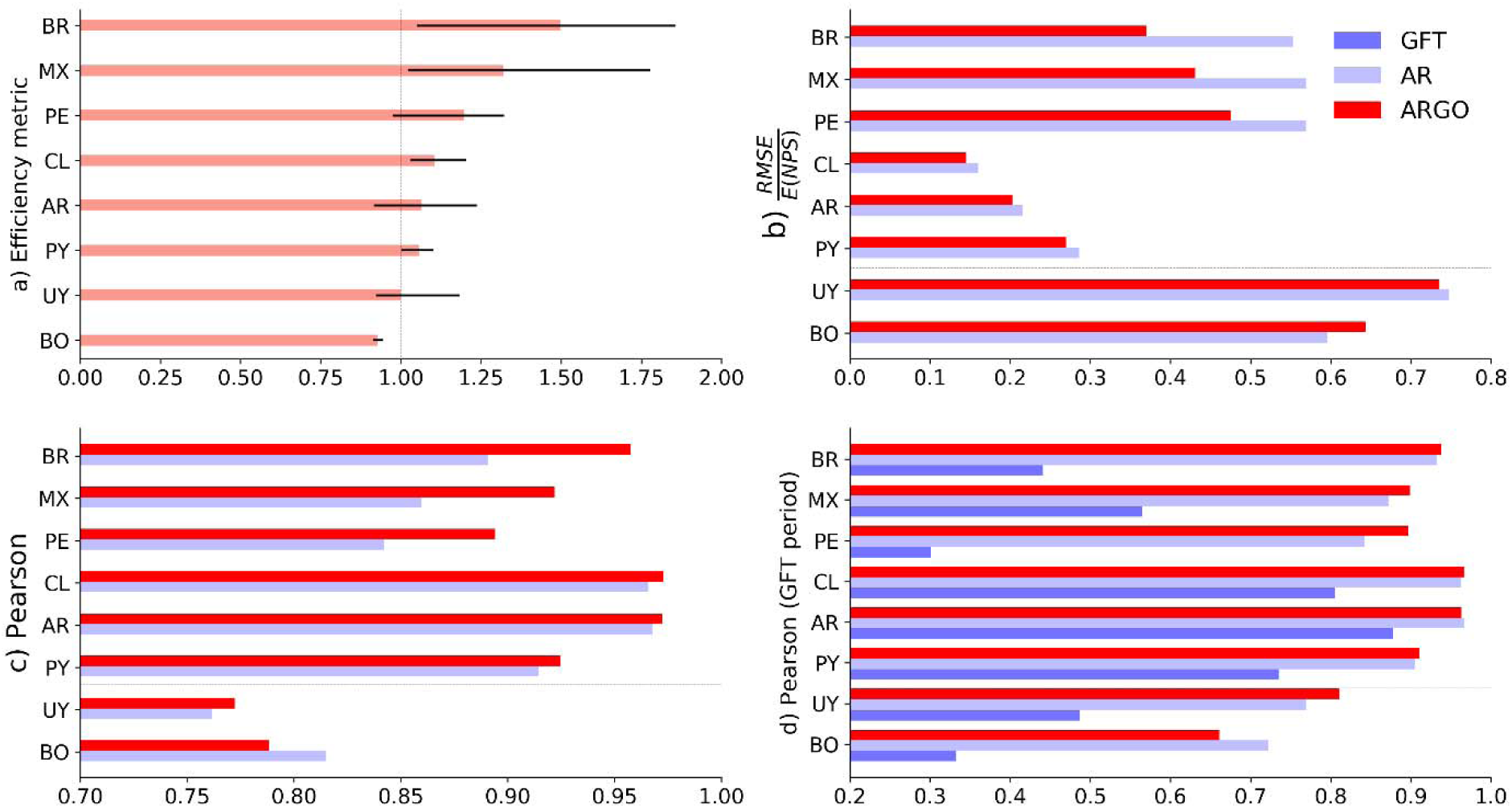
Set of bar graphs that show our performance metrics to assess predictive power of ARGO. (a) Efficiency metric values (salmon color) for each individual country with their respective 90% interval (solid black line). (b) Root mean square error values for ARGO (red) and AR52 (light blue) during the whole study period. Each country’s RMSE value is normalized by their respective average NPS over the whole study period to avoid scale differences in visualization. c) Pearson correlation scores of ARGO and AR during the full period. d) Pearson correlation values for ARGO (red), AR (gray) and GFT (blue) during the period in our study where GFT was active. For more detailed information about metric scores, refer to Table 1 in the appendix.

ARGO displays improvement in 6 countries in terms of the efficiency metric, reaching significant error reductions in Brazil (155 to 104 or 33%), Mexico (243 to 184 or 24%), Peru (48 to 40 or 16%) and Chile (131 to 119 or 9%). ARGO consistently outperforms GFT on Pearson correlation during the time period when GFT was active in every country and improves upon AR in all countries except Bolivia and Uruguay over the whole study period, reaching significant correlation increases in Brazil (from 0.891 to 0.957), Mexico (from 0.86 to 0.92) and Peru (from 0.84 to 0.89). See Supplementary Materials.

## Discussion

ARGO’s prediction performance shows that Internet search volumes and historical flu activity, when combined with dynamic machine learning techniques, can effectively detect real-time suspected flu cases in several Latin American countries. Our results considerably outperform the historical predictive performance of Google Flu Trends by using Internet search trends in a robust, self-adjusting manner. The overall improvement of ARGO over the baseline autoregressive model indicates that Internet search engine data, even in middle income countries, provides increased responsiveness to changing disease trends. This improvement is clear in Brazil, Chile, Mexico, Peru, Paraguay, and Argentina, whereas in Uruguay and Bolivia, the inclusion of Google search data does not seem to improve the baseline model.

The availability of an online tool to select relevant flu-related terms (Google Correlate) that track historical flu activity was found to be a critical element for ARGO to improve performance over the autoregressive benchmark (Argentina, Chile, Mexico, Peru, and Brazil), suggesting that the most meaningful flu-related search queries are country-specific. In countries where many weekly data points were missing on FluNet, like Uruguay, ARGO’s predictive ability was reduced. Our best performance was seen in Brazil, Mexico, Peru, where flu data was collected consistently every week during this study’s time-period (See supplementary materials).

Based on our previous research findings monitoring Dengue and Zika activity in Latin America [20, 21], we chose the number of suspected influenza cases (as captured by the FluNet’s NPS) as our gold standard for our prediction tasks. Our choice was based on the intuitive fact that flurelated Google search activity is higher when more people “suspect” they may be affected by flu like symptoms, regardless of the outcome of any lab test. As such, our models may prove useful to improve the timely allocation of resources in healthcare facilities in situations when increased numbers of people, with flu-like symptoms and respiratory needs, may need to be seen. Our choice of gold standard is to be contrasted with a previous study conducted by Pollet, who chose the positive influenza proportion of processed specimens as their ground truth [19].

## Notes

**Funding** MS and FL were partially funded by the Centers for Disease Control and Prevention’s Cooperative 375 Agreement PPHF 11797-998G-15. LC was partially funded by a grant with register 734557 by the Mexico National Council of Science and Technology.

**Conflict of Interest** None

